# Human primary plaque cell cultures to study mechanisms of atherosclerosis

**DOI:** 10.1101/2023.02.09.527800

**Authors:** Michele F. Buono, Ernest Diez Benavente, Lotte Slenders, Daisey Methorst, Daniёlle Tessels, Eloi Mili, Roxy Finger, Daniek Kapteijn, Mark Daniels, Noortje A. M. van den Dungen, Jorg J. A. Calis, Barend M. Mol, Gert J. de Borst, Dominique P. V. de Kleijn, Gerard Pasterkamp, Hester M. den Ruijter, Michal Mokry

**Affiliations:** Laboratory of Experimental Cardiology, University Medical Center Utrecht, Utrecht, The Netherlands; Central Diagnostics Laboratory, University Medical Center Utrecht, Utrecht, The Netherlands; Department of Cardiology, University Medical Center Utrecht, Utrecht, The Netherlands; Center for Translational Immunology, University Medical Center Utrecht, Utrecht, The Netherlands; Pediatric Immunology & Rheumatology, Wilhelmina Children’s Hospital, University Medical Center Utrecht, Utrecht, The Netherlands; Department of Vascular Surgery, University Medical Centre Utrecht, Utrecht, The Netherlands

**Keywords:** Phenotypic switch, Smooth muscle cells, Plaque cells, Disease modelling

## Abstract

Plaque smooth muscle cells are critical players in the initiation and advancement of atherosclerotic disease. They produce extracellular matrix (ECM) components, which play a role in lesion progression and stabilization. Despite clear phenotypic differences between plaque smooth muscle cells and vascular smooth muscle cells (VSMCs), VSMCs are still widely used as a model system in atherosclerotic research.

Here we present a conditioned outgrowth method to isolate plaque smooth muscle cells. We obtained plaque cells from 27 donors (24 carotid and 3 femoral endarterectomies). We show that these cells keep their proliferative capacity for eight passages, are transcriptionally stable, retain donor-specific gene expression programs, and express extracellular matrix proteins (*FN1, COL1A1, DCN*) and smooth muscle cell markers (*ACTA2, MYH11, CNN1*).

Single-cell transcriptomics of plaque tissue and cultured cells reveals that cultured plaque cells closely resemble the myofibroblast fraction of plaque smooth muscle cells. Chromatin immunoprecipitation sequencing (ChIP-seq) shows the presence of histone H3 lysine 4 dimethylation (H3K4me2) at the *MYH11* promoter, pointing to their smooth muscle cell origin. Finally, we demonstrated that plaque cells can be efficiently transduced (>97%) and are capable to take up oxidized LDL (oxLDL) and undergo calcification.

In conclusion, we present a method to isolate and culture primary human plaque cells that retain plaque myofibroblast-like cells’ phenotypical and functional capabilities - making them a suitable *in vitro* model for studying selected mechanisms of atherosclerosis.

## 1 Introduction

Atherosclerosis is a complex systemic condition leading to plaque formation in the arterial walls. Plaque smooth muscle cells play a crucial role in this process. In atherosclerotic tissues, plaque smooth muscle cells are responsible for the production of ECM within the lesion (1), the formation of the fibrous cap (2) and they are likely involved in driving the differences in the cellular composition between plaque phenotypes (3–6).

Even though these cells are mostly derived from vascular smooth muscle cells (2) (VSMCs) they are functionally and phenotypically different. For example, a fraction of plaque smooth muscle cells showed to lack detectable expression of some canonical SMC genes, but express mesenchymal (Sca1, CD105), as well as myofibroblast (αSMA, PDGFRB) markers (7). Moreover, lineage tracing studies showed that part of plaque smooth muscle cells are derived from transdifferentiated cells with non-VSMC origins (7,8). Despite the clear phenotypic differences between VSMCs and plaque smooth muscle cells, VSMCs are still widely used as a model system in atherosclerotic research.

Several groups previously isolated and cultured plaque smooth muscle cells by mincing or enzymatically digesting atherosclerotic tissues (9–13). According to those studies, the isolated cells resembled smooth muscle cells in culture. However, the explant approach by mincing and culturing the pieces of tissues leads to highly heterogeneous cultures and has a low yield (9–11). The enzymatic digestion activates the expression of chemokines and adhesion molecules, leading to extensive leukocyte adhesion (12), affecting the yield and growth of cells derived from plaque (10,13).

Here we present a robust and reproducible outgrowth explant-based method for isolating plaque smooth muscle cells from human atherosclerotic tissues. The cell morphology and transcriptome closely resemble the myofibroblast fraction of plaque smooth muscle cells and the cells retain patient-specific gene expression profiles.

## 2 Results

### 2.1 Outgrowth cultures from fresh atherosclerotic plaques

We tested two culture media - IMDM-FBS (14) and HAM F12K Complete (designed in-house, see Methods), in combination with Matrigel, Vitronectin, and Fibronectin pre-coated plates, and in the presence and absence of Vitamin C. Figure 1 provides an overview of the methodology of the conditioned outgrowth method, which is based on the capability of plaque cells to migrate from 2-3 mm^2^ plaque pieces (Fig. S1 A) and proliferate in the dish. We compared the culture conditions by observing the cell’s morphology and growth using transmitted light microscopy (Fig. S1 B). We evaluated the combination of fibronectin coating, HAM F12K Complete and Vitamin C for the yield and morphology of the cells after 14 days of culture (Fig S1 B – n). The addition of Vitamin C was crucial to maintain the cell proliferation in culture (Fig S1 B – m, n), and has a limited effect on gene expression (Fig. S1 C).

**Figure 1.**
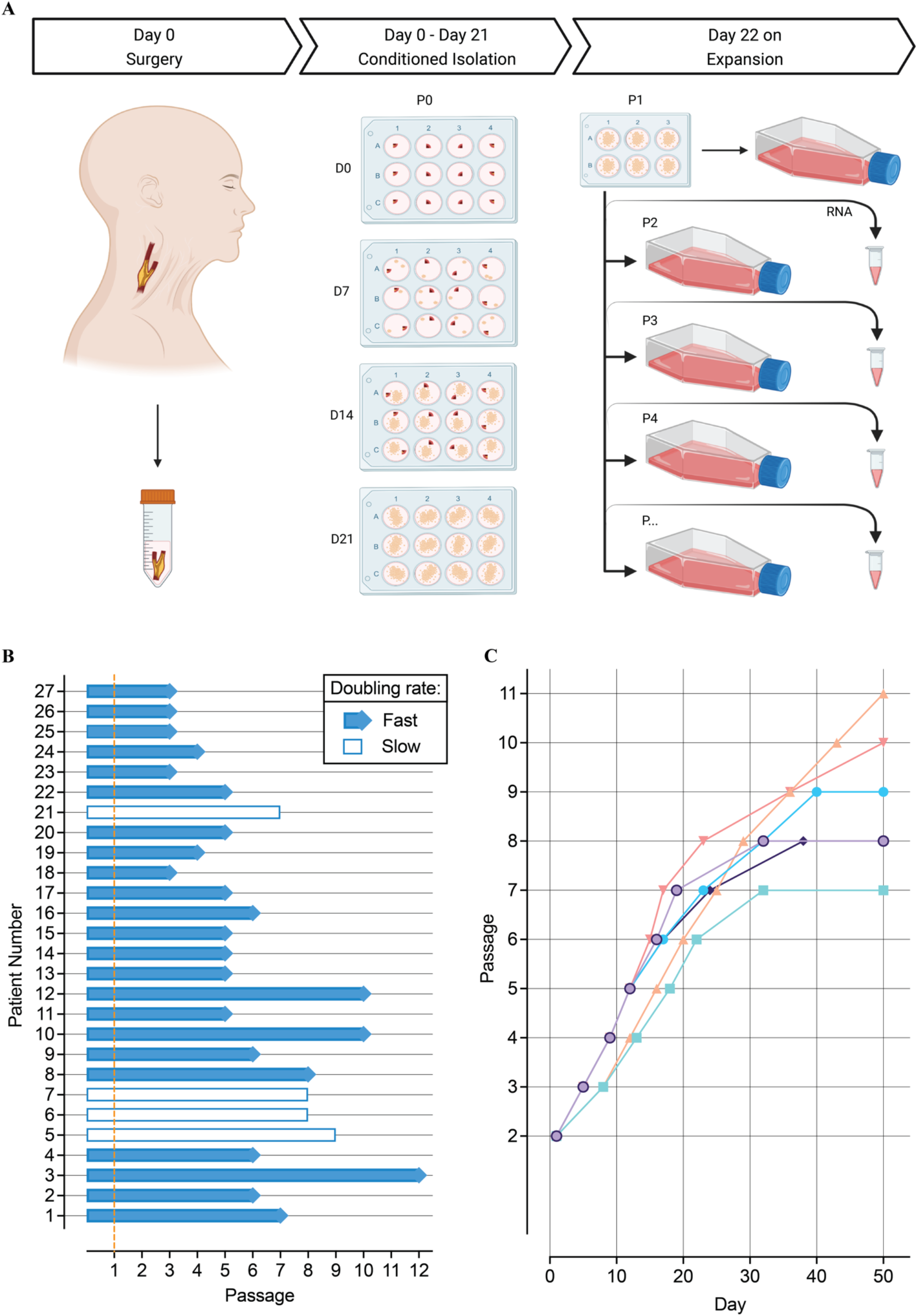
Isolation and growth of plaque cells. **(A)** Conditioned outgrowth method to isolate and expand primary plaque cells from fresh human atherosclerotic plaques obtained from endarterectomy patients. Plaques were obtained after surgery (D0), cut and placed in a pre-coated fibronectin 12 well plate until day 14 (D14), while refreshing the culture medium every other day. At D14, atherosclerotic pieces were removed from the dish and the cells migrated and clustered in colonies were left to grow until day 21 (D21). On day 21, the cell colonies were passed to a 6-well plate (p1) and subsequently expanded. From p2 on, plaque cells were sub-cultured until 70-80% confluency and sampled for mRNA extraction over the culture. **(B)** Growth status of individual plaque cell lines until their latest passage. The orange dashed line indicates the end of the isolation procedure. **(C)** Growth curves of 6 individual examples of plaque cells showing their proliferative capacity.

We successfully isolated and cultured human primary plaque smooth muscle cells from 27 out of 31 endarterectomy patients (Table 1). The isolation of 4 plaque cell lines failed due to fungal contaminations. From the 27 atherosclerotic plaques, 24 (11 males, 13 females) were obtained from the carotid and 3 (1 male, 2 females) from the femoral (Table S1) artery.

**Table 1.**
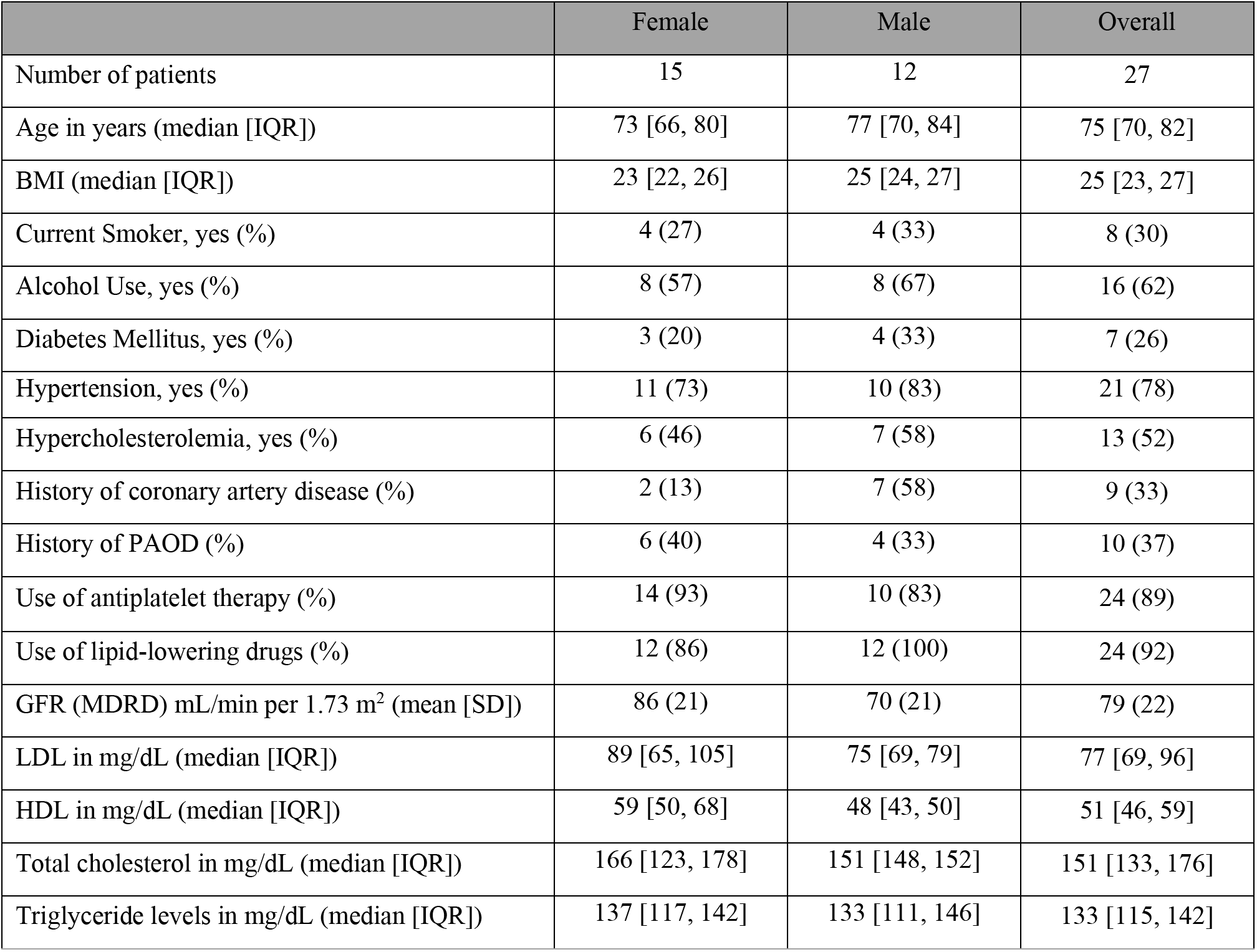
Baseline characteristics of patients used for the isolation of plaque cells.

We expanded all the isolated plaque cells for at least 3 passages (Fig. 1 B). The cells have an average doubling rate of approximately 4 days until passage 8, after that the proliferative capacity decreases (Fig. 1 C). Typically, 8 million plaque cells can be obtained from a single isolation by passage 5.

### 2.2 Characterizing plaque cells

To characterize the cells, we performed confocal microscopy, showing that cultured plaque cells were positive for α-SMA (Fig. 2 A, i - ii) and presented a stretched morphology when stained for phalloidin (F-Actin) (Fig. 2 A, iii - iiii). Next, we measured the expression of canonical VSMC markers such as ACTA2, MYOCD, MYH11, MYH10, PDGFB, KLF4, TPM4, MMP2, MMP2, CNN1 using qPCR in cultured plaque cells and human coronary artery smooth muscle cells (HCASMCs) (Table S1). Plaque cells had comparable expression levels of all the markers among donors, and they showed lower levels of MYOCD and PDGFb and higher ACTA2, MYH11, KLF4 and CNN1 levels when compared with HCASMCs (Fig. 2 B).

**Figure 2.**
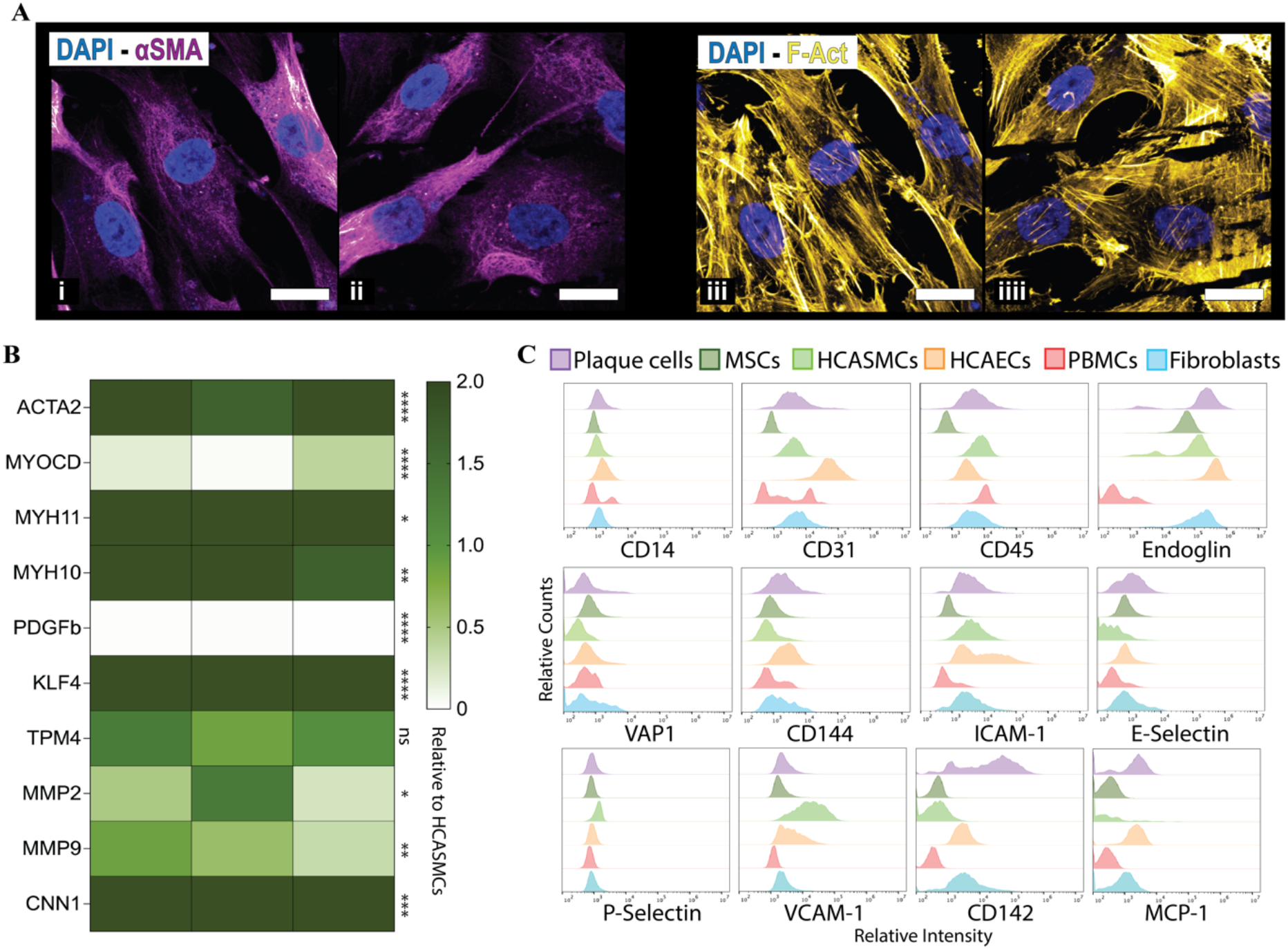
Characterization of plaque cells. **(A)** Representative confocal images of two plaque cell lines showing positivity for alpha-SM Actin (i – ii) and morphological tracts via phalloidin staining (F-Actin, iii - iiii). Scale bar 20 μm. **(B)** Gene expression analysis of 3 individual plaque cell donors for the canonical VSMC markers relative to HCASMCs. Each column represents one patient and the corresponding gene expression pattern. * p ≤ 0.05, ** p ≤ 0.01, *** p ≤ 0.001, **** p ≤ 0.0001. **(C)** Flow cytometry analysis comparing expression profiles of plaque cells to other cell types involved in atherogenesis. The histograms show the “Relative Counts” on the Y axis and the “Relative Intensity” on the X axis.

Flow cytometry was used to compare the protein expression profile of various cell markers in plaque cells and other cell types involved in atherogenesis or present in artery walls (15) such as HCASMCs, human coronary artery endothelial cells (HCAECs), fibroblasts, mesenchymal stem cells (MSCs) and peripheral blood mononuclear cells (PBMCs). Accordingly, isolated plaque cells best matched with VSMCs and fibroblasts profiles but with different expression of markers like E-Selectin, CD142 and MCP1 (Fig. 2 C).

### 2.3 Transcriptional stability of plaque cells in culture

To understand whether the plaque cells maintain a stable transcriptome or undergo transcriptional changes in culture, the RNA was extracted from plaque cells of 12 individual donors (6 males, 6 females), from passage 2 to 5, and was studied by RNA-seq (Table S1). Principal component analysis (PCA) suggests that plaque cells of the same donor, at different passages, group together. This demonstrated that the donor characteristics were retained in culture and had higher impact on the cell transcriptome than prolonged culture (Fig. 3 A).

**Figure 3.**
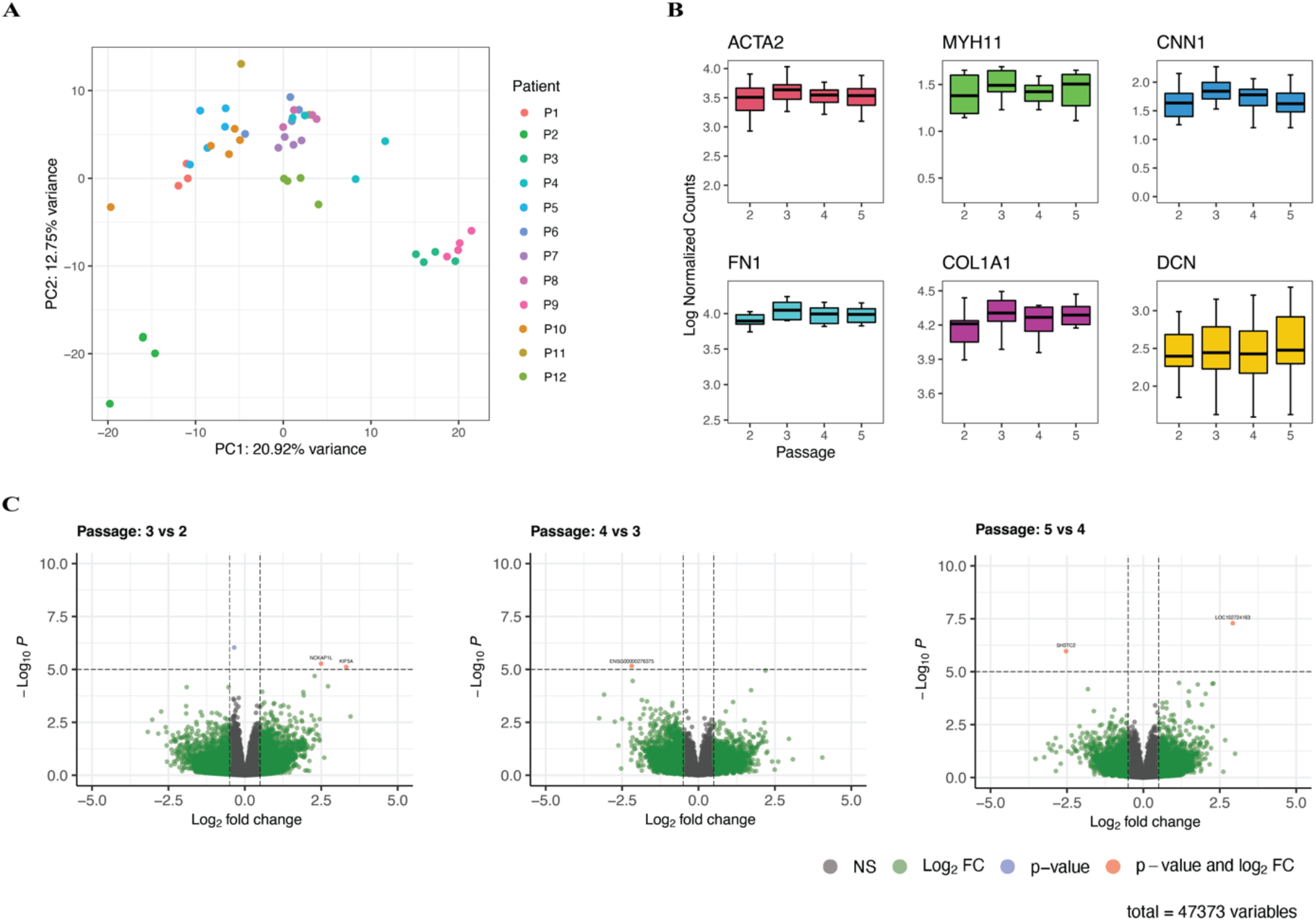
Stable transcriptome maintained over passages. **(A)** Principal component analysis, based on the top 500 variable genes, of plaque cells from 12 patients. **(B)** Box plots showing the expression of canonical SMC and fibroblast genes, such as ACTA2, MYH11, CNN1, FN1, COL1A1, DCN, throughout passages (from 2 to 5). **(C)** Differential gene expression analysis between passage 2, 3, 4 and 5.

The expression of canonical SMC and fibroblast markers (ACTA2, MYH11, CNN1, FN1, COL1A1, DCN) (16) was consistent from passage (p) 2 to 5, indicating transcriptome stability (Fig. 3 B). Using linear models, only 12 genes showed a time-trend across these passages (Fig. S2 C). In addition, we performed differential gene expression between consecutive passages. This showed that only 3 coding genes (NCKAP1L, KIF5A, SH3TC2) and 2 non-coding ones were differentially expressed in independent comparisons from p2 to p5, suggesting that plaque cell expression profiles were stable over the passages (Fig 3 C). Differential gene expression analysis of p1 versus other passages highlighted several immune-cell specific genes (e.g. CD74, CD4) (Fig. S2 E) and enrichment analysis revealed ongoing inflammatory processes (Fig. S2 F). In line with this, p1 is grouped separately from the higher passages in the PCA plot (Fig. S2 D). These findings indicated that plaque cell cultures at p1 potentially still contained some cells expressing leukocyte markers, which were strongly reduced from p2 onwards.

Plaque cells derived from TEA patients clustered differently compared to CEA patients (Fig. S2 A). Differential gene expression analysis showed transcriptional differences between the cells isolated from carotid and femoral plaques. The TEA-derived cells overexpressed specific HOX-class genes such as HOXC9 (Fig. S2 B), which is known for playing a role in the development and rostro-caudal gene expression patterns (17). These findings resonate with findings in dogs where transcription patterns of vascular cells were dependent on vessel localization (18).

### 2.4 Transcriptome analysis points to the shared smooth muscle cell and fibroblast characteristics of isolated plaque cells

We projected the RNA-seq data derived from the pooled plaque cells from the 12 individual patients onto several existing single-cell RNA sequencing (scRNA-seq) dataset. The first was generated in our laboratory from 46 human carotid plaques (26 males and 20 females, Fig. 4 A) (27). The isolated plaque cells most closely resemble ACTA2^+^ SMC transcriptomic profiles (Fig. 4 A). We found that they best matched with SMC1 cluster (Fig. 4 B), representing cells with smooth muscle cell and fibroblast characteristics and hence defined as myofibroblasts.

**Figure 4.**
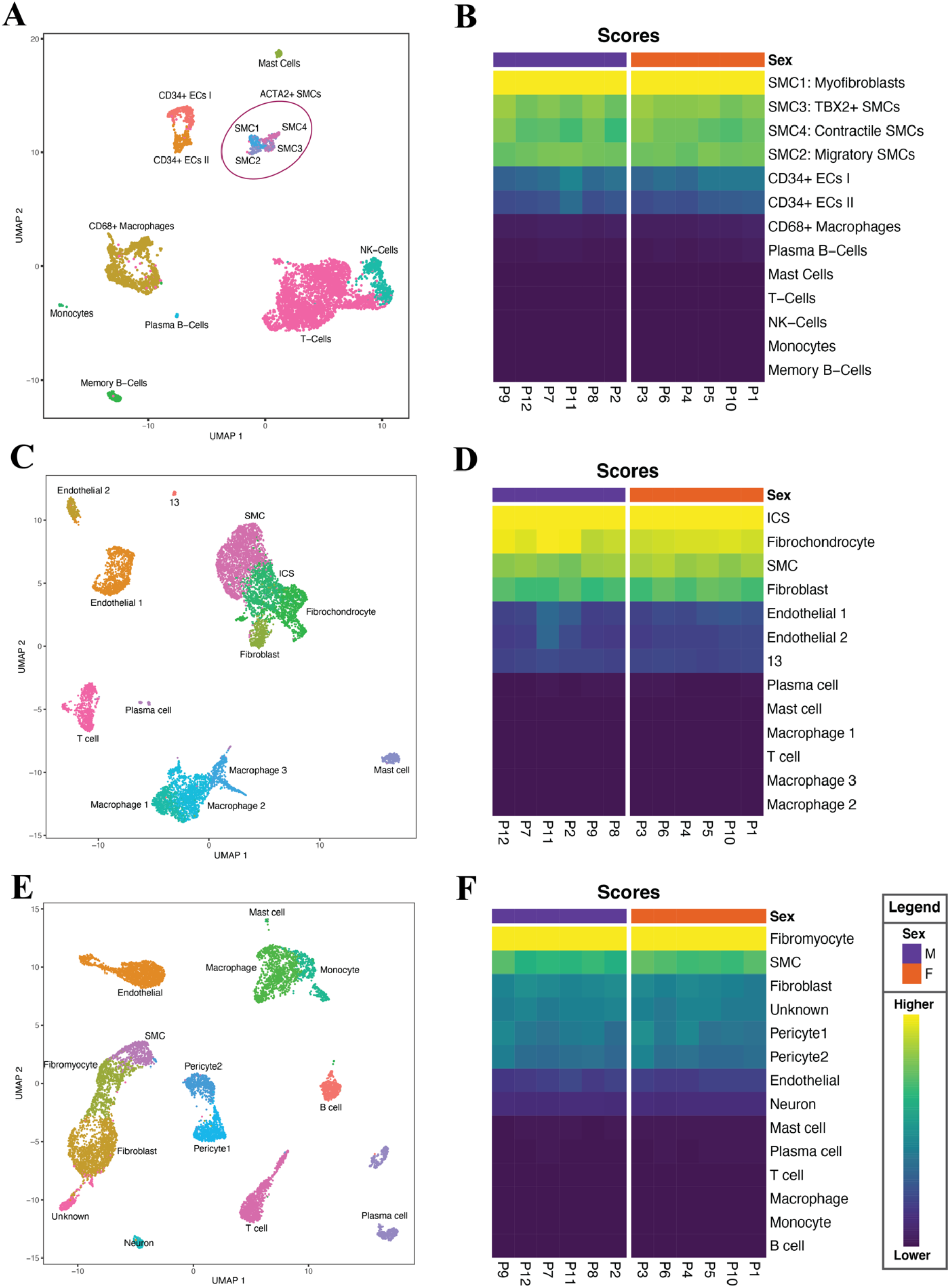
Transcriptome characterization of plaque cells. **(A)** UMAP showing different cells clusters identified in the scRNA-seq of 46 atherosclerotic plaques of carotid endarterectomies donors, previously generated by our group (19). **(B)** Heatmap showing that plaque cells are matched with the ACTA2+ VSMCs cluster, and specifically with SMC1 subcluster which represent cells with myofibroblast characteristics. **(C)** UMAP of a scRNA-seq of 3 atherosclerotic plaques of carotid endarterectomies donors, generated and published by Pan et al. (20). **(D)** Heatmap showing that plaque cells matched with ICSs which according to the author is a cluster of cells representing an intermediate state between SMCs and fibroblasts/fibrochondrocytes. **(E)** UMAP of scRNA-seq of 4 atherosclerotic coronary plaques, generated and published by Wirka et al. (5). **(F)** Heatmap showing that plaque cells matched with the fibromyocyte cluster which represent cells with SMC origins and FB characteristics.

To validate our finding, we used another human scRNA-seq carotid plaque dataset generated by Pan et al. (Fig. 4 C) (20). This showed that plaque cells mostly match the intermediate cell state (ICS) cluster (Fig. 4 D). This cluster represents an intermediate cell state between SMCs and fibroblasts/fibrochondrocytes. We then projected our data on a scRNA-seq dataset of human atherosclerotic coronary arteries (n=4 patients) generated by Wirka et al. (Fig. 4 E) (5). Interestingly, our plaque cells matched fibromyocytes (Fig. 4 F) which represent modulated cells with a SMC origin and fibroblast characteristics. Although plaque cells matched cell clusters named differently among datasets, generated from diverse diseased vascular beds, they all showed the characteristics of an intermediate phenotype between smooth muscle cells and fibroblasts.

### 2.5 Single-cell RNA sequencing revealed phenotypic gradient of plaque cells

Next, given the diversity of plaque smooth muscle cells within atherosclerotic plaques (21), we performed scRNAseq on individual plaque cells to understand the heterogeneity of the isolated cells. We used a SPLiT-seq method with passage-synchronized plaque cells from 4 individual patients (2 females and 2 males respectively; p4) to compare these with the scRNA-seq dataset of carotid plaque tissue.

A total 1834 cells remained after QC, of which 469 for patient 5, 686 for patient 6, 379 for patient 7, and 300 for patient 8. This data was then projected to the scRNA-seq dataset from carotid plaques using singleR. Plaque cells were distributed amongst the four SMC subclusters with the majority matching SMC1 (Fig. 5 A). On average plaque cells match 88.6% SMC1, 9.9% SMC3 1.2% SMC4 and 0.3% SMC2 (Fig. 5 B).

**Figure 5.**
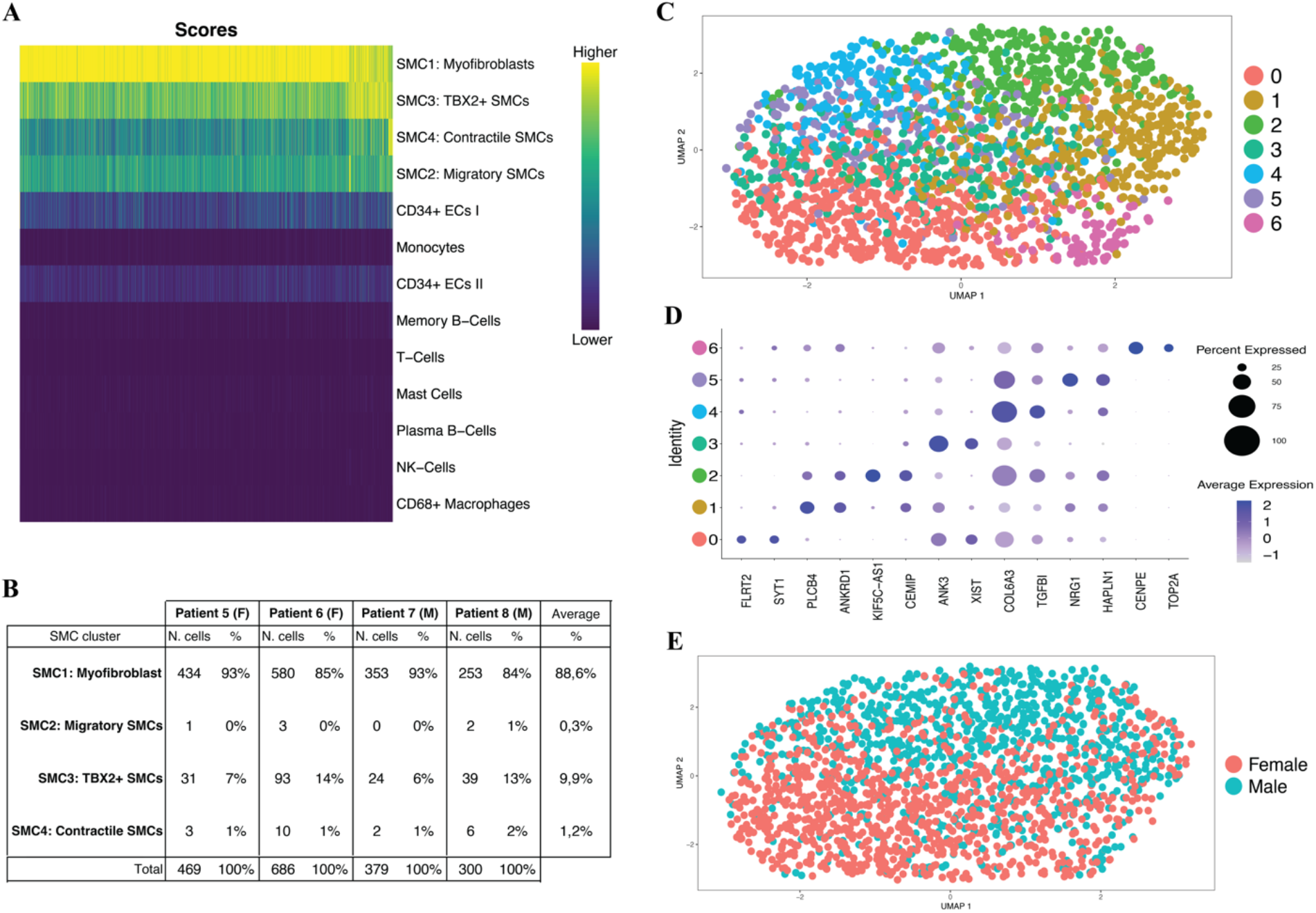
Different subsets of plaque cells. **(A)** Heatmap showing that plaque cells matched with the four SMC clusters, mainly with SMC1. **(B)** Table showing the distribution of plaque cells from 4 individual patients within the four SMC clusters. **(C)** UMAP generated by plotting plaque cells with similar gene expression profiles, showing that seven different clusters of plaque cells can be identified**. (D)** The 2 top genes expressed in the 7 clusters were plotted in a Dot plot showing differences among plaque cell cluster profiles. **(E)** UMAP of male and female plaque cells showing sex-specific patterns.

We clustered the cells using Seurat and used UMAP to project the individual cells based on their expression similarities. We identified seven clusters (Fig. 5 C – D) which were not clearly separated in the UMAP projection (Fig 5 D). Interestingly, considering the sexes of the donors, male and female cells were separated in UMAP projection (Fig. 5 E). We found the XIST as one of the genes correlating with this clustering (Fig. 5 D). Furthermore, sex differential gene expression showed that the sex-chromosome expression complement of the donor is maintained *in vitro* (Table S2, Table S3).

### 2.6 Plaque cells origin

To study the origin of plaque cells, we utilized SMC-specific epigenetic modification – H3K4me2 on MYH11 promoter, previously used to trace cells that originate from SMCs. (22,23). We performed chromatin immunoprecipitation sequencing (ChIP-seq) for the H3K4me2 and H3K27Ac (Fig. 6). In both HCASMCs and plaque cells, H3K4me2 showed an increased signal compared to the background on the promoter of MYH11. Based on this finding, the dimethylation mark found in this specific locus of MYH11 promoter suggests that plaque cells have SMCs origin. Moreover, H3K27Ac showed a lower signal in plaque cells when compared with HCASMCs, suggesting that MYH11 promoter is less active in plaque cells.

**Figure 6.**
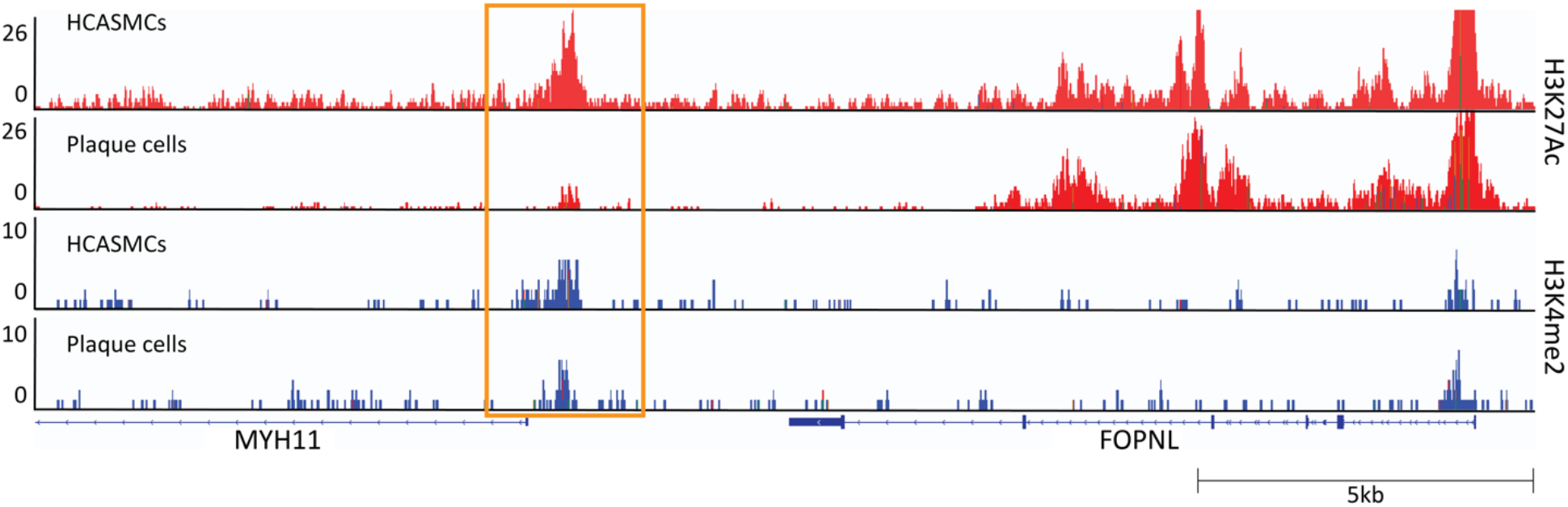
Plaque cells have SMC origin. ChIP-seq occupancy of H3K4me3 and H3K27ac in HCASMCs and plaque cells at a specific locus of MYH11 promoter.

### 2.7 Plaque cells as a cellular model for atherosclerosis

Transduction efficiency can be a limiting factor in the usability of a cellular models. Therefore, to investigate the transferability of plaque cells, we transduced plaque cells with CVM-GFP lentiviruses and showed that most of the cells were GFP-positive (Fig. 7 E). Transduction efficiency was measured via flow cytometry demonstrating that > 97% of the cells were expressing GFP and hence successfully transduced (Fig. 7 F).

**Figure 7.**
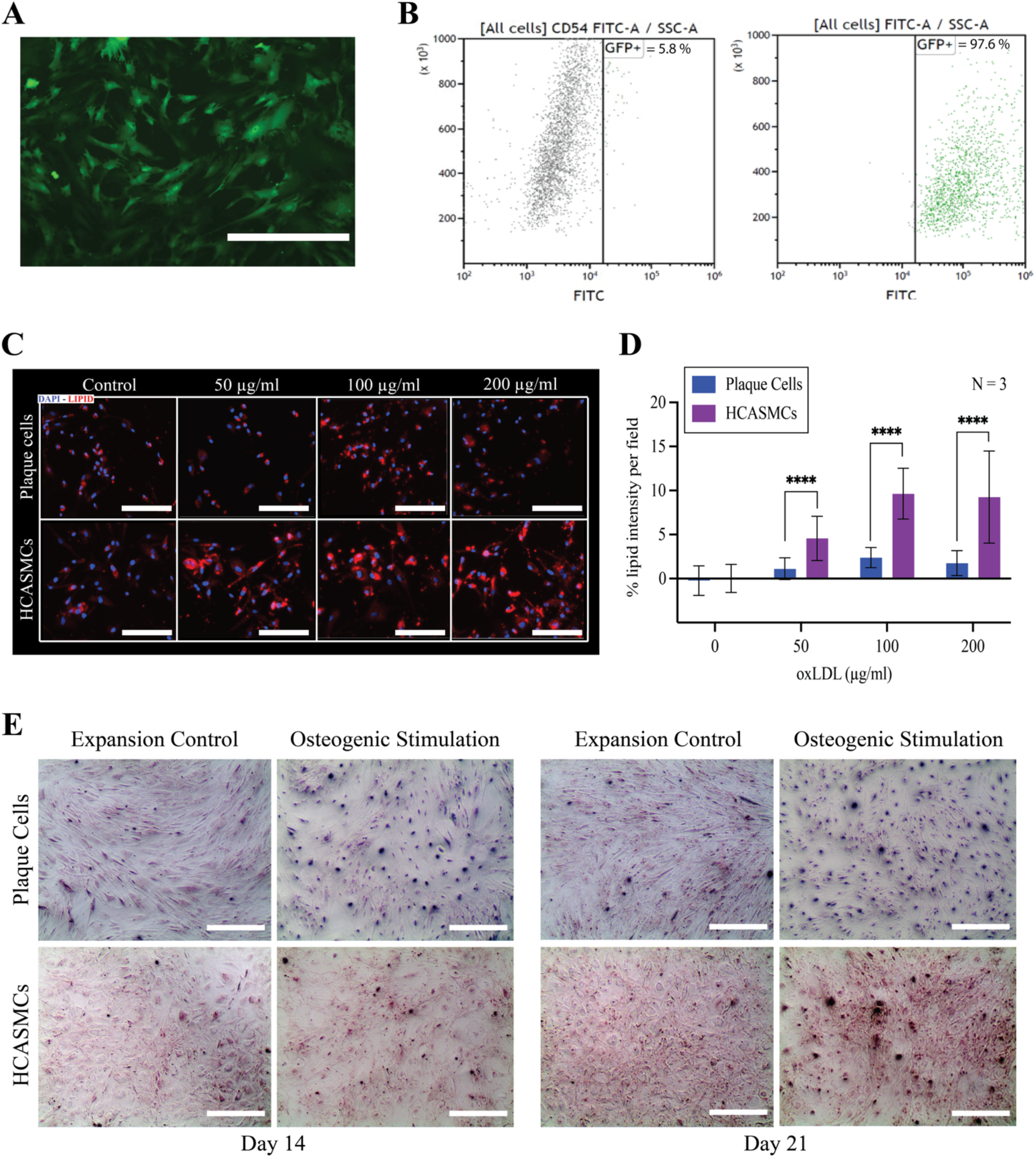
Application of plaque cells. **(A)** Representative fluorescence image showing positive-GFP plaque cells 72 hours after lentiviral transduction. Scale bar = 400 μm. **(B)** Flow cytometry scatter plots showing GFP-positive plaque cells 72 after transduction. Control non-transduced plaque cells are shown in (i) while the transduced ones are in (ii). **(C)** Representative fluorescence images of plaque cells and HASMCs, showing the uptake of oxLDL (red) after 24 hours exposure at different concentrations (0, 50, 100, 200 μg/ml). Scale bar = 200 μm. **(D)** Barplot showing the quantified oxLDL taken up by plaque cells and HCASMCs when compared with non-treated control (0). Normalized for confluency. **** p ≤ 0.0001. **(E)** Representative transmitted light images showing Alazarin red staining to visualize the deposition of calcified matrix in plaque cells and HASMCs after 14 and 21 days of osteogenic stimulation and expansion control culture. Scale bar = 400 μm

Then, we used plaque cells as a cellular model for the most important SMC-related mechanisms involved in the progression of atherosclerosis - oxLDL uptake and calcification. Specifically, we focused on the accumulation of oxidized LDL (oxLDL), relevant for the formation of the lipid core (24), and calcified matrix leading to vascular calcification and to severe events such as occlusion of the lumen, plaque rapture, thrombus formation and infraction (25).

To check if plaque cells were capable of uptake oxLDL and if there was any difference with commercial SMCs, we exposed plaque cells and HCASMCs to different oxLDL concentrations, as 50, 100, 200 μg/ml. After 24 hours of exposure, nuclei and lipids were stained, and internalized oxLDL was quantified. We showed that both plaques cells and HCASMCs taken up oxLDL at all concentrations (Fig. 7 A). However, plaque cells accumulate 4folds less oxLDL than HCASMCs (p ≤ 0.0001; Fig. 7 B).

Then, to study the capabilities of plaque cells undergo osteogenic transformations, we osteogenically stimulated plaque cells and HCASMCs for 21 days. After, we stained our cells with Alizarin red, which binds to the calcium of calcified matrix. We found the deposition of calcified matrix after 14 days of culture in plaque cells while we observed less deposition in HCASMCs. After 21 days of osteogenic stimulation both cell types showed similar deposition of calcified matrix. (Fig. 7 C).

## 3 Discussion

In this study, we established an outgrowth method based on the capability of plaque smooth muscle cells to migrate and outgrow from the dense structure of atherosclerotic tissue to the culture dish. We reliable and effectively obtained and cultured plaque cells from 27 donors, indicating high reproducibility of the isolation protocol. The yield of plaque cells and their proliferative activity *in vitro* is lower when compared with commercial HCASMCs. This is unsurprising, considering that in vivo plaque cells are constrained in the dense atherosclerotic tissue which limits proliferation and the supply of essential nutrition and growth factors. Indeed, atherosclerotic plaques grow over the years before becoming symptomatic or getting diagnosed (26). However, 8 x 10^6^ plaque cells can be obtained by p5 with a sustained doubling rate of 4 days until passage 8. The possibility to keep them in culture for weeks, makes plaque cells suitable for the development of several functional tests.

Our transcriptomic studies highlighted that plaque cells retain donor characteristics, and these are maintained during prolonged culture. Plaque cells have a unique expression profile, expressing markers of both plaque SMCs and fibroblasts, which better resemble myofibroblasts. They express ACTA2, TAGLN, CNN1, MYH11, TPM2 and IL6, FBLN1, DCN, which are accepted as SMC and fibroblast markers, respectively (16). The downregulation or absence of MYOCD is considered as the marker that indicates the switch of medial VSMCs toward this more plastic phenotype (21). Also, plaque cells showed a higher expression of ECM markers and adhesion protein such as fibronectin (FN1), which suggest their role as the key determinant of plaque structure as connective tissue promote the formation of the protective fibrous cap (27,28).

Moreover, differential gene expression indicated that plaque cells maintain a stable transcriptome over the culture but also that cells at p1 retain inflammatory signals (CD4, CD74), may be due leukocyte infiltration. However, from p2 onwards, the cultures lose these signals and become more homogenous. High-throughput scRNA-seq showed the presence of 7 subclusters of plaque cells which have slightly different expression profiles which can be related to their plasticity. While, considering the sex of the donors at single cell level, female and male cells cluster separately, demonstrating further that donor characteristics are maintained *in vitro*.

Furthermore, ChIP-seq showed H3K4me2 mark in the MYH11 promoter confirming the SMC origin of plaque cells. This finding, together with the retained patient-specific gene expression, makes plaque cells suitable for the *in vitro* characterization of cellular mechanisms involved in plaque progression and for studying functional consequences of atherosclerosis SMC-specific key driver genes.

During plaque formation and development *in vivo*, SMCs lose or acquire certain markers, transdifferentiating to different cell types influenced by outer stimuli (29–31). However, their response may be modulated by the density and the structure of the surrounding connective tissue. Thus, *in vitro* studies using plaque cells may prove useful in exploring their response to external signals and assist in the study of the molecular mechanisms of formation and progression of atherosclerotic lesions. In light of this, we showed that plaque cells are capable of taking up oxidized LDL. This would be useful for future studies to understand how SM-derived foam cells originate which may help on improve our understanding of sex-specific differences in atherosclerotic plaques. Indeed, plaque cells from males and females may uptake oxLDL differently, which may drive the process by which women develop plaques richer in ECM components and with a smaller or absent lipid/necrotic core when compared to men. Moreover, the presence of calcification within atherosclerotic plaques suggest the similar to chondrocytes, osteocytes, and even osteoclasts (32). Calcification is considered a marker of plaque vulnerability and hence more prevalent in plaques developed by men than women (33,34). This way, to prove the potential of plaque cells and understanding whether they are a key player in determining plaque phenotypes, we performed a calcification study showing that they were capable to calcify. Furthermore, we also demonstrated that plaque cells can easily be transduced, which will open a plethora of new possibilities for further functional tests like gene silencing or editing studies, of significant therapeutic value in the seek for new gene therapies and druggable targets.

In conclusion, we are aware of all the limitations of cultured cells, especially in that they may differ from the original *in vivo* phenotype. However, we demonstrated that plaque cells retain patient-specific gene expression, are of SMC-origin and express crucial myofibroblast markers, which may inform our knowledge of the original tissue. Cultured plaque cells could be used to better study mechanisms affecting plaque formation and progression in a sex-specific manner but also to search for the new markers of plaque vulnerability. Moreover, plaque cells may be a good model for *in vitro* modulation of plaque cell death, regulation of interleukin and adhesion molecule expression, and testing of anti-atherosclerotic drugs.

## 5 Materials and Methods

### 5.1 Conditioned outgrow isolation and culture of human primary plaque cells

Primary plaque cells were isolated from fresh atherosclerotic plaque tissues obtained from patients underwent carotid or thromb- (femoral) endarterectomies (CEA or TEA). Informed consent was obtained from all patients preoperatively. Fresh tissues were collected in Hank’s Balanced Salt Solution 1X (HBSS, cat. 14025-50, Gibco) on ice within 30 min of operative resection. Tissue fragments were then dissected into 2-3 mm^3^ pieces. One or two pieces were placed in each well of a twelve-well plate, pre-coated with 2 μg/cm^2^ Fibronectin (cat. F1141, Sigma Aldrich). The pieces were cultured for fourtheen days, the first seven days in HAM F12K Complete containing antimicrobial agent for primary cells (Primocin, cat. ant-pm2, InvivoGen) (1:500) and then in complete medium only. The plaque pieces were removed from the wells at day 14 and cell colonies were kept in culture. At day 21, the colonies were dissociated and subcultured. Plaque cells were cultured by replenishing complete medium one other day and subcultured when 70-80% confluence at the density of 5 × 10^5^ cells per T75 dish. Cell counts were determined using the TC20 (Bio-Rad Laboratories, Hercules, CA, USA).

### 5.2 HAM F12K Complete Medium

Complete medium contains: HAM’s F12K (Kaighn’s) Nut mix 1x (cat. 21127022, Gibco) supplemented with 10% (v/v) Heat Inactivated FCS (Corning), 1% (v/v) Penicillin-Streptomycin (10,000 U/ml, cat. 15140122, Gibco), 1% (v/v) Insulin Transferrin Sodium Selenite solution (ITS, cat. 41400045, Fisher Scientific), 10 mM HEPES (cat. 15630056, Fisher Scientific), 10 mM TES (cat. T1375, Sigma Aldrich), 30μg/ml Endothelial Cell Growth Factors (ECGS, cat. 02-102, Merck/Sigma Aldrich), 2.5μg/ml of Vitamin C solution (L-Ascorbioc Acid, cat. A4544, Sigma Aldrich). Vitamin C solution were added weekly freshly made upon use.

### 5.3 FCS-depletion Medium

It consists of basal medium HAM’s F12K (Kaighn’s) Nut mix 1x supplemented with 2% (v/v) Insulin Transferrin Sodium Selenite solution, 20 mM HEPES, 20 mM TES, 60 μg/ml Endothelial Cell Growth Factors, 5 μg/ml of Vitamin C solution (added weekly and freshly made upon use).

### 5.4 qPCR

Along the cultures, samples of 3 x 10^5^ plaque cells were collected, lysed and stored at −80°C in 350 μl of lysis buffer (RA1, cat. 750961, Macherey-Nagel) every passaging step. Total RNA was isolated according to the supplier’s protocol (Nucleospin RNA, Macherey-Nagel). A detailed summary of the mRNA samples per plaque cell lines can be found in Table S1.

Transcription of 300 ng of DNA-free RNA into cDNA was performed using the qScript cDNA Synthesis Kit (Quantabio, #95047). qRT-PCR was performed using iQ SYBR Green Supermix (Bio-Rad) with specific primers in a CFX96 Touch Real-Time PCR detection system (Bio-Rad): 5 min at 95°C, followed by 40 cycles of 15 s at 95°C, 30 s at specific annealing temperature, and 45 s at 72°C, followed by melting curve analysis to confirm single product amplification. Messenger RNA (mRNA) expression levels were normalized to human heterochromatin protein 1 binding protein 3 (hHP1BP3) reference gene mRNA expression (ΔCt). Relative differences were calculated (ΔΔCt) and presented as fold induction (2^-ΔΔCt^). Primers used are shown in the Table S4. Data from three different plaque cell lines were used to assess up/down regulation of canonical SMC markers when compared with commercially available human coronary artery smooth muscle cells (HCASMCs).

### 5.5 Generation of Growth Curves

Cells were subcultured when 70-80% confluence at the density of 5 × 10^5^ cells per 75 cm^2^ and used from passage 2 until they showed a significant decrease in proliferation. The number of days between two subculturing procedures was annotated during the cultures. Data were plotted in GraphPad Prism (V9).

### 5.6 Flow Cytometry

Dissociated cells were directly transferred to flow cytometry tubes (BD Biosciences) and stained with a live/dead discriminator using 1:1000 Zombie Nir Dye in PBS. Then, the samples were permeabilized using a buffer containing 20% FBS, 3% BSA, 0.2% Saponin in PBS for 10 min at RT and stained for CD14, CD31, CD45, Endoglin, VAP1, CD144, ICAM-1, E-Selectin, P-Selectin, VCAM-1, CD142, MCP1 (Table S2) in 1% BSA, 0.1% Saponin in PBS in the dark at RT for 1 hour. After staining, the samples were resuspended in in FACS buffer, containing 2% FBS, 5mM EDTA, 0.01% Sodium Azide in PBS and measured by recording 3 × 10^4^ events using CytoFLEX and Gallios cytometers (Beckman Coulter). The resulting data were analysed using FlowJo v10 (TreeStar). Antibodies used are shown in the Table S5.

### 5.7 RNA sequencing

RNA library preparation was performed, adapting the CEL-Seq2 protocol for library preparation (35,36). The initial reverse-transcription reaction primer was designed as follows: an anchored polyT, a unique 6bp barcode, a unique molecular identifier (UMI) of 6bp, the 5’ Illumina adapter and a T7 promoter. Complementary DNA was used for vitro transcription reaction (AM1334; Thermo-Fisher). The resulting amplified RNA (aRNA) was fragmented and cleaned. RNA yield and quality were checked by Bioanalyzer (Agilent).

cDNA library construction was initiated according to the manufacturer’s protocol, with the addition of randomhexRT primer as random primer. PCR amplification was performed with Phusion High-Fidelity PCR Master Mix with HF buffer (NEB, MA, USA) and a unique indexed RNA PCR primer (Illumina) per reaction. Library cDNA yield was checked by Qubit fluorometric quantification (Thermo-Fisher) and quality by Bioanalyzer (Agilent). Libraries were sequenced on the Illumina Nextseq500 platform with paired end, 2 × 75 bp (Utrecht Sequencing Facility).

Upon sequencing, fastq files were de-barcoded and split for forward and reverse reads The reads were demultiplexed and aligned to human cDNA reference (Ensembl version 84) using the BWA (0.7.13) by calling ‘bwa aln’ with settings -B 6 -q 0 -n 0.00 -k 2 -l 200 -t 6 for R1 and -B 0 -q 0 - n 0.04 -k 2 -l 200 -t 6 for R2, ‘bwa sampe’ with settings -n 100 -N 100. Multiple reads mapping to the same gene with the same unique molecular identifier (UMI, 6bp long) were counted as a single read.

### 5.8 Single cells RNA sequencing (SPLiT-seq)

Passage synchronized plaque cells were collected at 70-80% confluency, counted using the TC20 (Bio-Rad Laboratories, Hercules, CA, USA) and centrifuged at 350 g for 6 min. The supernatant was removed, and the pellet was rinsed in 1 ml of PBS pH 7.4 (Gibco). At this stage the cell suspensions were fixed and processed using the SPLiT-seq library preparation method. Briefly, cells were fixed with 1% formaldehyde, permeabilized and counted. Then the cells divided in 8000 cells per well together with reverse transcription (RT) mix (Maxima H minus Reverse Transcriptase ThermoFisher) and incubated for RT barcoding. Then the cells were pooled, and divided into DNA barcode plate 1, together with ligation mix and incubated for 30 min at 37°C. After, blocking solution was added to each well and again incubated for 30min at 37°C. The cells were then collected from plate 1, pooled with addition of extra ligase (T4 DNA ligase NEB), divided over wells in DNA barcode plate 2 and incubated for 30 min at 37°C. Blocking solution was added to each well and again incubated for 30min at 37°C. Then cells were pooled, lysed and washed. Sublibraries were made and purified for cDNA with dynabeads+streptavidine, after TemplateSwitch the cDNA was amplified (with 2xKapa HiFi hotstart Mastermix Kapa biosystems) and size selected via ampure. The cDNA is tagmented and amplicons/libraries are generated with Illumina Nextera XT library prep kit, the libraries are bioanalyzed and sequenced. The SPLiT-seq protocol was adapted from the study of Rosenberg AB et al. (37) to include a sample-specific first barcode for multiplexing. In SPLiT-seq, reverse transcription (RT) and barcode ligation steps occur in fixed intact cells. Cells are pooled and split before the barcoding steps, such that different barcode combinations are introduced to individual cells. Reads with the same barcode combination are analyzed together to determine which transcripts an individual cell expressed.

### 5.9 Chromatin Immunoprecipitation sequencing (ChIP-seq)

Chromatin immunoprecipitation was performed using the MAGnify Chromatin Immunoprecipitation System (Thermo Fisher Scientific, 492024) as stated in the manufacturer’s manual. In short, the Dynabeads were incubated with the antibodies, while in the meantime crosslinking of the cells was performed with 1% formaldehyde. The reaction was stopped with 1.25 M glycine. Cells were lysed and sonicated with the S2 sonicator (Covaris) to shear the chromatin into 100-300 bp fragments following the program stated in the protocol. The microTUBE AFA Fiber Crimp-Cap 6×16mm (Covaris, SKU: 520052) was used with the sonicator. 10 μl of the diluted chromatin was set aside for the input control sample. The diluted chromatin and the Dynabeads were added together, and several washing steps were performed followed by protein digestion using proteinase K. The crosslinking was reversed at 65 °C and the samples were purified using the ChIP DNA Clean and Concentrator kit (Zymo Research, D5205). The DynaMagTM-2 Magnet (Thermo Fisher Scientific, 12321D) was used when working with the magnetic beads.

The immunoprecipitated DNA were used for creating libraries which could then be sequenced. The libraries were created with the NEXTFLEX Rapid DNASeq Kit 2.0 (PerkinElmer, NOVA-5188-01) and the NEXTFLEX-HT Barcodes (PerkinElmer, NOVA-5188-12). The DNA in the libraries were visualized using the FlashGelTM DNA Cassette (Lonza, 57032) together with the Gel Loading Buffer II (Invitrogen, AM8546G) and the 50 bp – 1.5 Kb FlashGel™ DNA Marker as ladder (Lonza, 57033). The 2100 Bioanalyzer of Automated Droplet Generator (Agilent) was used to visualize the fragment sizes in each sample. The Qubit^®^ 3.0 Fluorometer (Thermo Fisher Scientific, Q33216) together with the QubitTM dsDNA HS Assay kit (Thermo Fisher Scientific, Q32854) was used to quantify the DNA concentration of each library. The libraries were pooled and send to the Utrecht Sequencing Facility (USEQ). The first run was analyzed using the Illumina NextSeq500 platform with a 1 x 75 bp High Output Run type. The last two runs were analyzed using the Illumina NextSeq2000 with a 2 x 50 bp Run type. Then, the mapped Next Generation Sequencing data against the human genome (GRCh37/hg19) was visualized using the Integrative Genomics Viewer (IGV).

### 5.10 oxLDL Uptake

Plaque cells and HASMCs were seeded (1 × 10^4^ / well) into a black 96 well plate and maintained in FCS-depletion medium for 24 hours. After 24 hours the medium was refreshed and oxidized lipoproteins (oxLDL, L34357, Thermo Fisher Scientific, 2.5 mg/ml) were added to the cultures. The oxLDL were diluted to 50, 100 and 200 in FCS-depletion medium and incubate at 37 C, 5% CO2 for the subsequential 24 hours. After, plaque cells were rinsed with 1X PBS (Gibco) and fixed for 15 min with 4% paraformaldehyde (PFA, Klinipath). Then, they were permeabilizated with 0.1% Triton X-100 in PBS for 15 minutes on a shaker at RT. After two washing steps with PBS, the cells were incubated with LipidSpot610 (biotium, 70069) to stain lipids for 30 minutes on a shaker at RT in the dark. Then, were washed with PBS and incubated with DAPI (1:10000) for 10 minutes on a shaker at RT in the dark. After, the cells were washed with PBS and imaged using a widefield microscope with CY5 2.0 filter for lipid visualization (Invitrogen, AMEP4956). The control conditions, cells without any exposure of oxLDL, were used as threshold for the blue (DAPI) and red (lipids) imaging parameters. The oxLDL taken up by the cells were quantified with ImageJ using a macro run protocol. Lipid intensities were multiplied for 50, to normalize the values obtained from ImageJ for the cell confluency (50%).

### 5.11 Osteogenic stimulation

Osteogenic stimulation of plaque cells an HASMCs was performed in 12-well plates, seeded in two replicate wells per donor in their expansion medium. Osteogenic stimulation was induced 24 hours post-seeding using DMEM supplemented with 10% FCS, 1% Glutamax and 1% penicillin/streptomycin, 50 μM L-Ascorbic acid 2-phosphate sesquimagnesium salt hydrate, 10 mM β-Glycerophosphate disodium salt hydrate and 0.1 μM water soluble dexamethasone (the last three all Sigma-Aldrich) (25). The cells were stimulated for 21 days, with osteogenic medium exchange every 3 to 4 days. Alizarin red staining was performed at day 14 and day 21, on the cells cultured with expansion control medium or osteogenic medium, to determine deposition of calcified matrix. Briefly, the cells were fixed for 15 min in 4% PFA, then each well was stained with 500 μl 0.5% Alizarin red (Sigma; in ddH2O, pH 4.2) for 10 min at room temperature and washed three times with ddH2O to remove unbound dye. Plates were allowed to dry, and staining was documented microscopically using a EVOS FL Cell Imaging System.

### 5.12 Transduction

Lentiviral vector containing GFP reporter gene, phage2-GFP (Addgene #86684), to stably express fluorescent human protein in mammalian cells was propagated in NEB Stable Competent E. coli (New England Biolabs). HEK293T/17 (ATCC CRL-11268) were maintained in antibiotic-free DMEM supplemented with 10% fetal bovine serum (v/v). On day one, 3 x 10^5^ cells per well, resuspended in 4 ml of medium, were seeded in a 6 well-plate. The following day, the antibiotic-free medium was replaced, 2 ml per well, two hours prior transfection. Later, the cells in each well were transfected with 1 μg of phage2GFP plasmid, 0.5 μg of pVSV-G envelope plasmid, 1 μg of pCMV (plasmid encoding viral packaging proteins) and 5 μl of P3000 diluted in 100 μL Opti-MEM with 7 μL of NaCl. Three days after transfection, medium was centrifuged at 500 g for 5 min to remove cell debris following filtration using 0.45 mm syringe filter. The CMV-GFP lentiviruses were further concentrated by ultracentrifugation with 20% (w/v) sucrose cushion.

Plaque cells were plated for transduction in 6-well plates (Corning) at the density of 1 × 10^5^ cells per well and maintained in HAM F12K Complete medium. After 24 h, CMV-GFP lentiviruses were added directly to the culture media in each well. 72 hours post-transduction, transduction efficiency was checked via imaging and flow cytometry was performed to confirm transduction.

### 5.13 Statistical and Data analyses

Graphical representation and statistical analysis of qPCR and oxLDL uptake data were obtained using GraphPad Prism V9. Unpaired t-test was used to compare the experimental groups with the control groups. Heatmap data are presented as mean of three replicates, while barplot data are presented as mean ± standard deviation (SD). A p-value ≤ 0.05 was deemed statistically significant.

Read counts obtained from RNA sequencing data were normalized and differential gene expression between groups was performed using DESeq2 in R. The screening criteria for DEGs were P-value < 0.00001 and Log2 fold change ≥ 0.5 and ≤ (−0.5). To generate the volcano plot “EnhancedVolcano” package was utilized. The genes significantly over-expressed are represented by the red dots on the left while the ones significantly under-regulated are represented by the red dots on the right. Pathway enrichment analysis were performed using differentially upregulated genes, the package “EnrichR” enriched for “GO_Biological_Process_2021” was used.

The SingleR function was run at default settings, using the log-normalized counts as input.

## 6 Supplementary Materials

**Figure S1.**
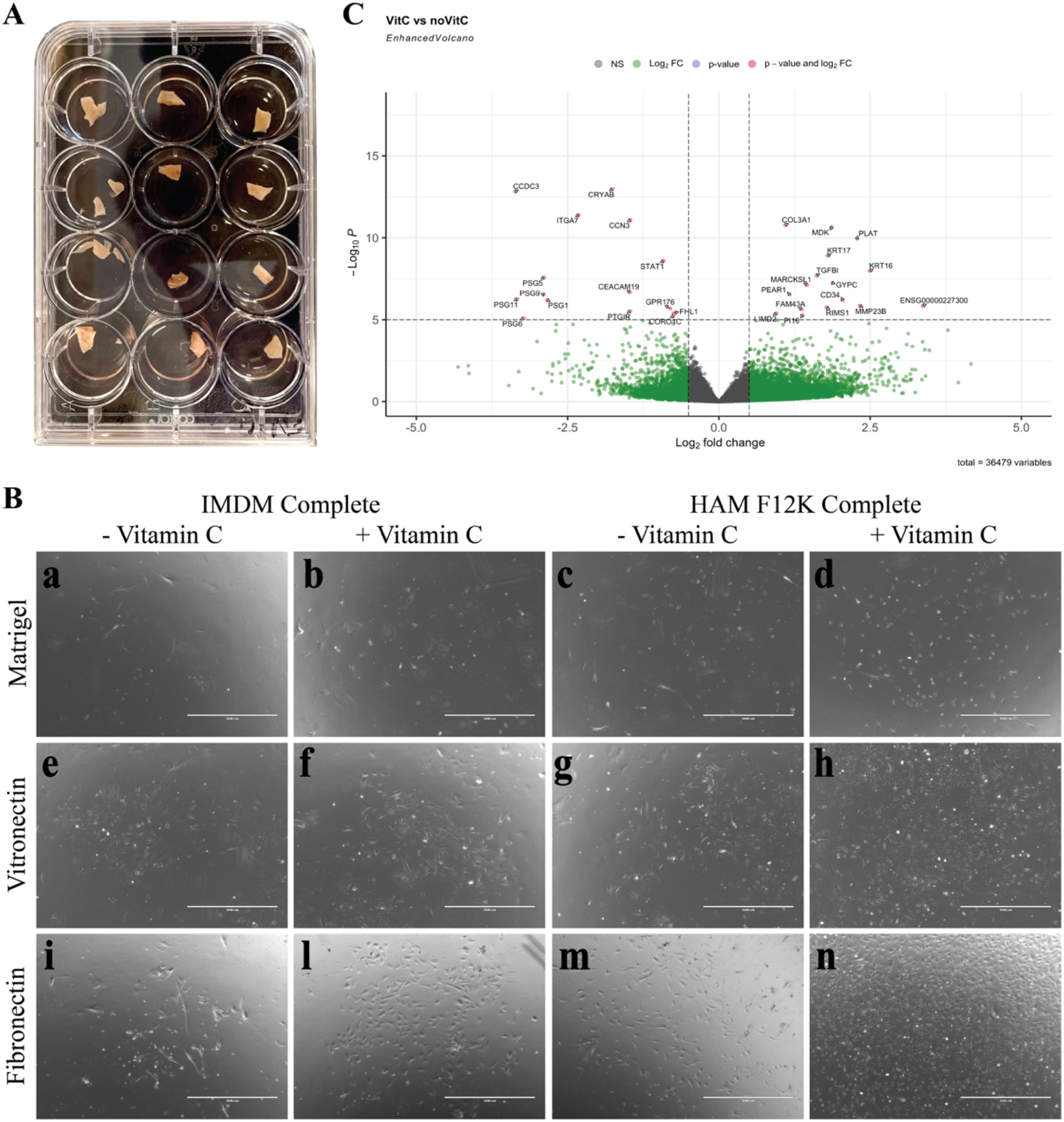
Establishment of the isolation method. **(A)** Representative picture of a 12 well-plate containing the 2-3 mm^2^ plaque pieces during cell isolation. **(B)** Representative transmitted light images taken after keeping the plaque pieces in culture for 14 days. Two different culture media were tested (IMDM-FCS and HAM F12K Complete), with and without the addition of Vitamin C, in combination with three coating materials (Matrigel, Vitronectin and Fibronectin). **(C)** Volcano Plot showing differentially expressed gene in plaque cells cultured in HAM F12K with and without Vitamin C.

**Figure S2.**
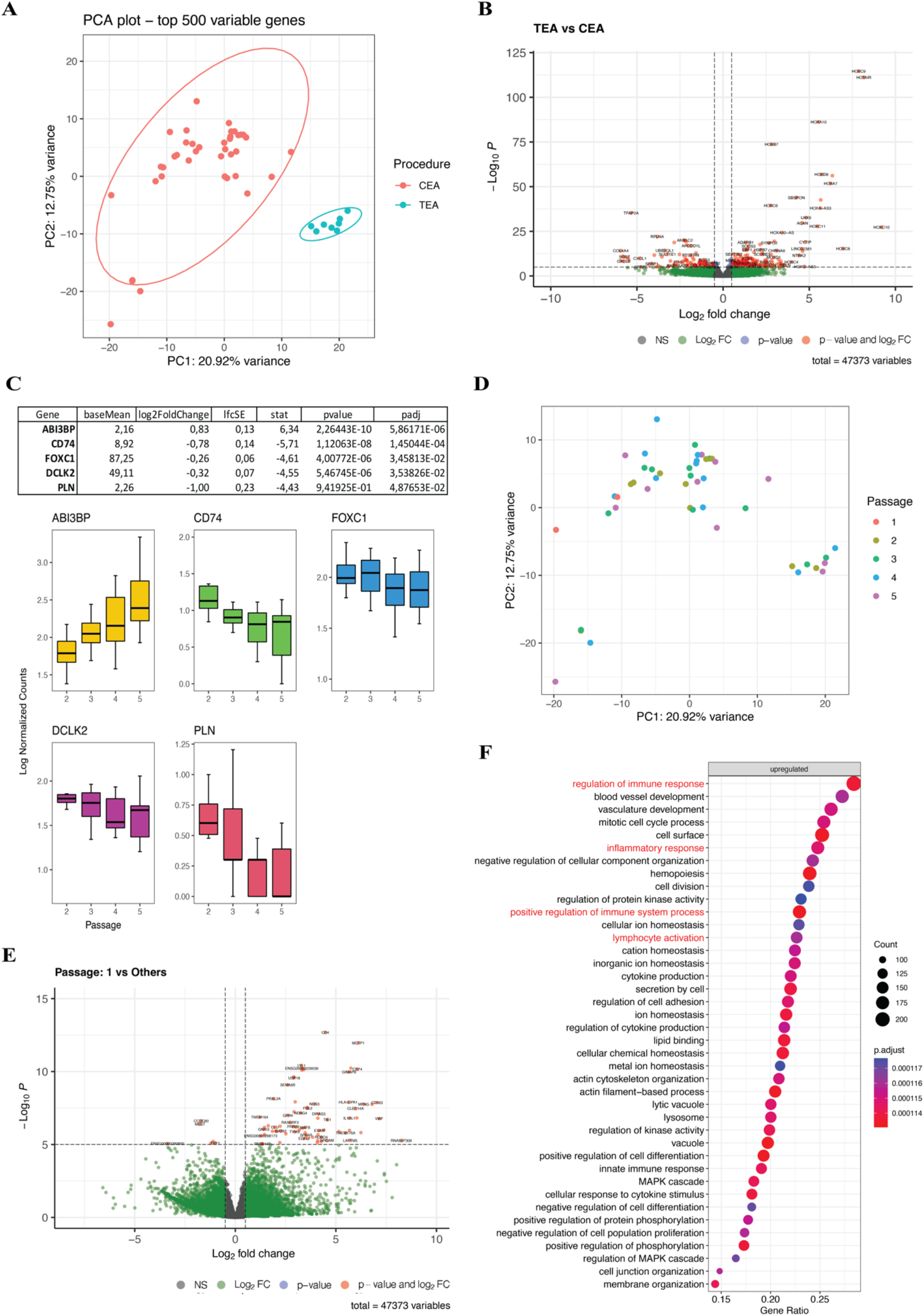
Transcriptional differences in plaque cells. **(A)** PCA plot of TEA- and CEA derived plaque cells. **(B)** Differential gene expression analysis of TEA vs CEA plaque cells. **(C)** Twelve time-trend genes h identified performing DEG analysis among passages. **(D)** PCA plot showing that passage 1 groups far from all the other passages. **(E)** Passage 1 showed several differential expressed genes, immune cells specific, when compared to the other passages. **(F)** Pathways analysis done for the genes differential expressed at passage 1.

**Table S1.**
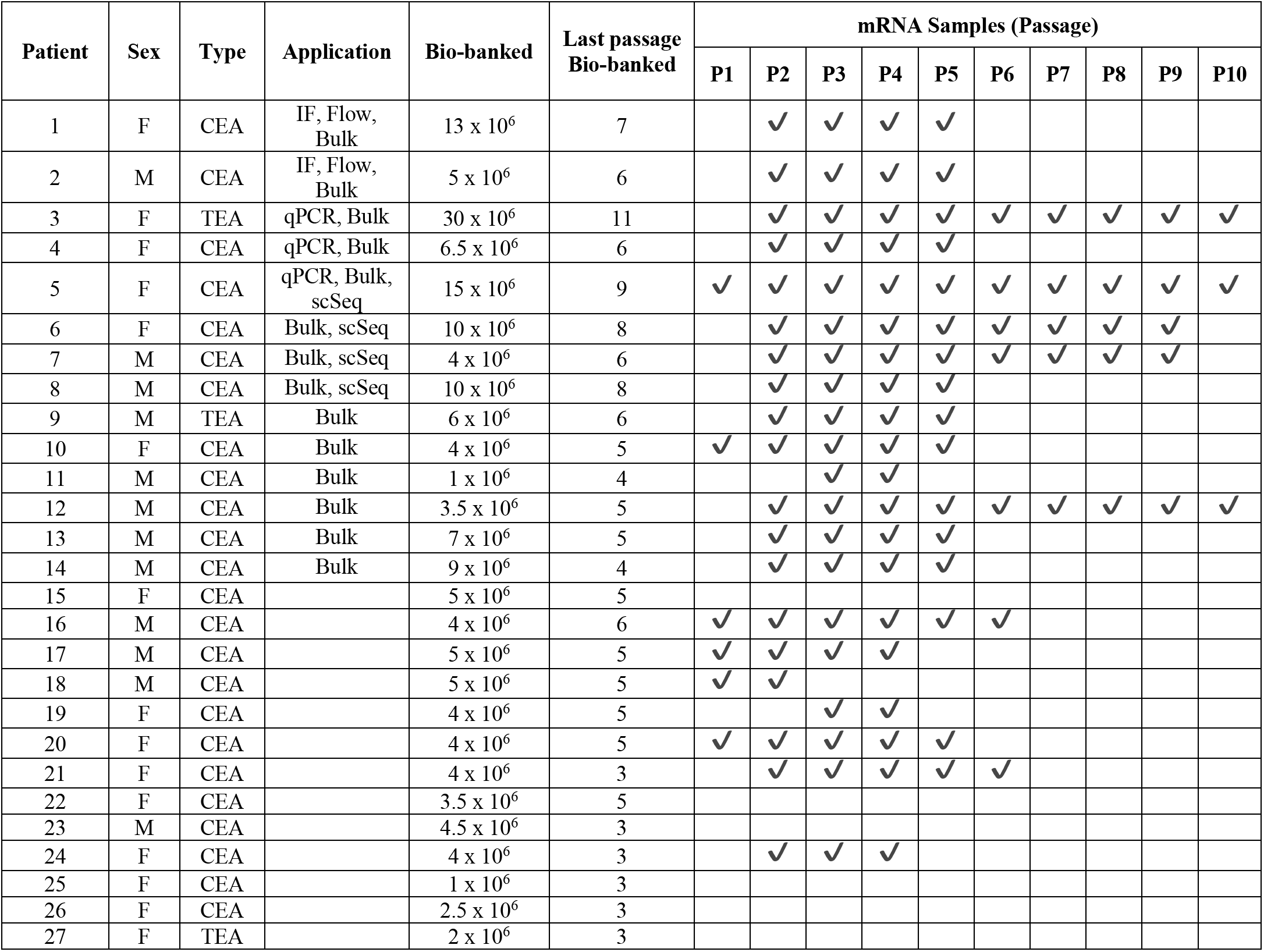
Patients used for plaque cell isolation, cells bio-banked and mRNA samples isolated along the cultures.

**Table S2.**
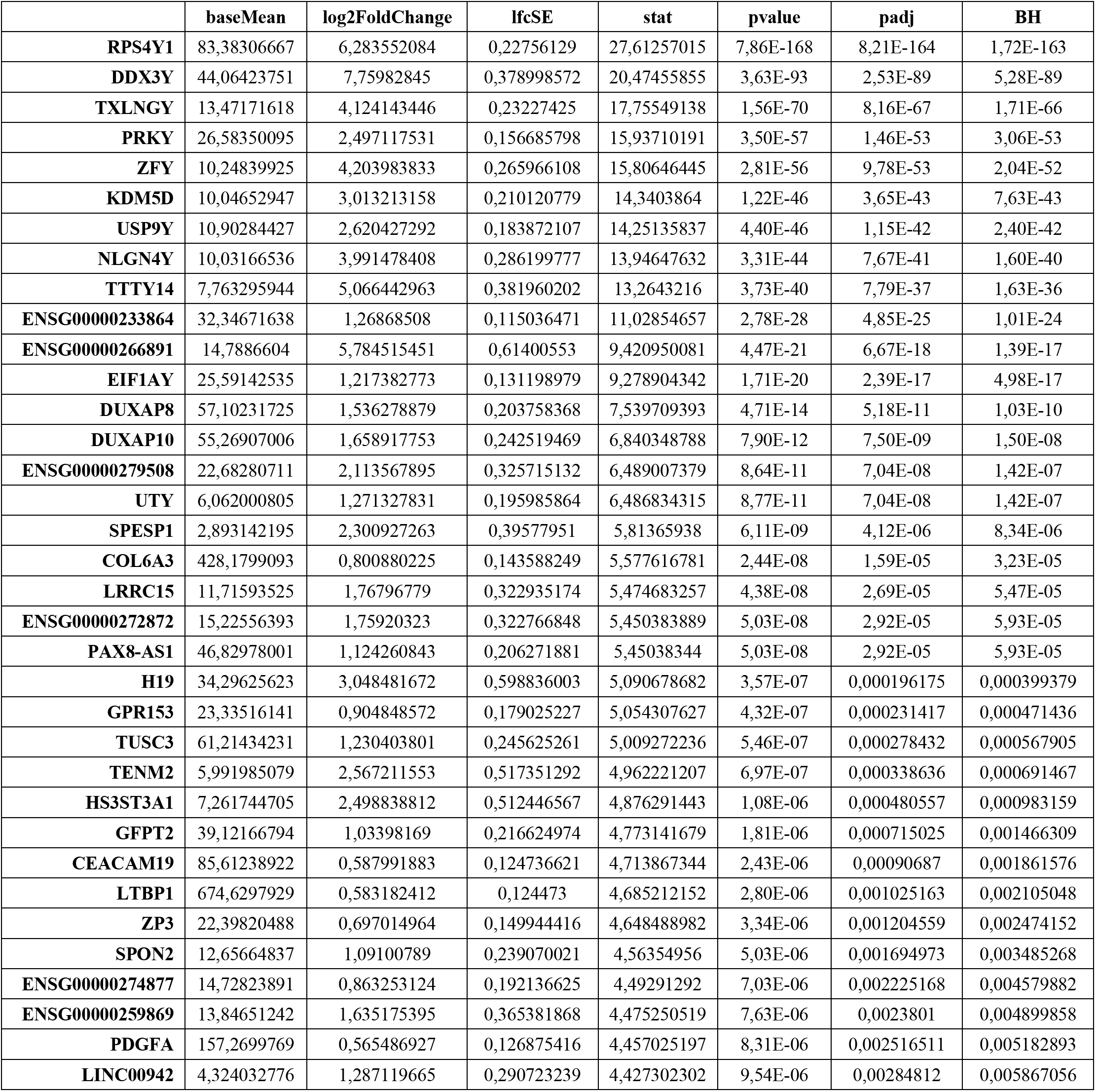
Overexpressed genes in male plaque cells after sex differential gene expression.

**Table S3.** Overexpressed genes in female plaque cells after sex differential gene expression.

|  | **baseMean** | **log2FoldChange** | **lfcSE** | **stat** | **pvalue** | **padj** | **BH** |
| --- | --- | --- | --- | --- | --- | --- | --- |
| **XIST** | 111,966434 | -5,635394 | 0,18118504 | -31,102976 | 2,20E-212 | 4,59E-208 | 9,58E-208 |
| **KDM5C** | 98,9524396 | -0,6099857 | 0,05142784 | -11,861 | 1,89E-32 | 3,58E-29 | 7,49E-29 |
| **PUDP** | 31,9847779 | -1,0863492 | 0,12559102 | -8,649895 | 5,15E-18 | 6,73E-15 | 1,41E-14 |
| **TSIX** | 148,523819 | -0,7501758 | 0,09184684 | -8,1676822 | 3,14E-16 | 3,86E-13 | 8,07E-13 |
| **OFD1** | 32,907764 | -0,782331 | 0,09827168 | -7,9608998 | 1,71E-15 | 1,98E-12 | 3,92E-12 |
| **RPS4X** | 1274,02182 | -0,5656642 | 0,07561463 | -7,4808838 | 7,38E-14 | 7,71E-11 | 1,53E-10 |
| **ZRSR2** | 25,2651395 | -0,7639271 | 0,1107962 | -6,8948849 | 5,39E-12 | 5,36E-09 | 1,07E-08 |
| **DHRS9** | 45,4017514 | -2,2295294 | 0,32799564 | -6,7974361 | 1,06E-11 | 9,67E-09 | 1,94E-08 |
| **GEMIN8** | 10,547866 | -0,8556854 | 0,1465671 | -5,8381817 | 5,28E-09 | 3,70E-06 | 7,49E-06 |
| **CA5BP1** | 78,6613873 | -0,5906688 | 0,10119579 | -5,8368908 | 5,32E-09 | 3,70E-06 | 7,49E-06 |
| **SUCNR1** | 17,3236946 | -2,7980761 | 0,50241966 | -5,569201 | 2,56E-08 | 1,62E-05 | 3,29E-05 |
| **TRPC4AP** | 76,7868846 | -0,5252915 | 0,10439897 | -5,0315776 | 4,86E-07 | 0,0002541 | 0,00051797 |
| **EHD3** | 60,9610731 | -0,6023846 | 0,12067191 | -4,9919207 | 5,98E-07 | 0,0002974 | 0,00060694 |
| **RTN1** | 21,46786 | -2,7666473 | 0,56612966 | -4,8869498 | 1,02E-06 | 0,0004755 | 0,00097192 |
| **IL33** | 7,39444364 | -2,4147178 | 0,50292321 | -4,8013648 | 1,58E-06 | 0,00068597 | 0,00140401 |
| **LIPA** | 267,460358 | -0,5197837 | 0,10835207 | -4,7971738 | 1,61E-06 | 0,00068618 | 0,00140502 |
| **RPS4XP6** | 17,6981069 | -0,5854776 | 0,1222421 | -4,7894924 | 1,67E-06 | 0,00069666 | 0,00142761 |
| **GSTT2** | 6,3718031 | -1,5386258 | 0,32177965 | -4,781613 | 1,74E-06 | 0,00069872 | 0,00143236 |
| **NES** | 66,0243542 | -1,1343093 | 0,2388671 | -4,7487048 | 2,05E-06 | 0,00079213 | 0,00162498 |
| **KIAA1549L** | 26,5581362 | -0,7413372 | 0,16024708 | -4,6262134 | 3,72E-06 | 0,00131884 | 0,00270967 |
| **RPS4XP11** | 50,5848409 | -0,5044443 | 0,10941216 | -4,6104955 | 4,02E-06 | 0,00137596 | 0,00282856 |
| **SLC16A14** | 7,48290265 | -3,6815184 | 0,81441713 | -4,5204334 | 6,17E-06 | 0,00204672 | 0,00420961 |

**Table S4.**
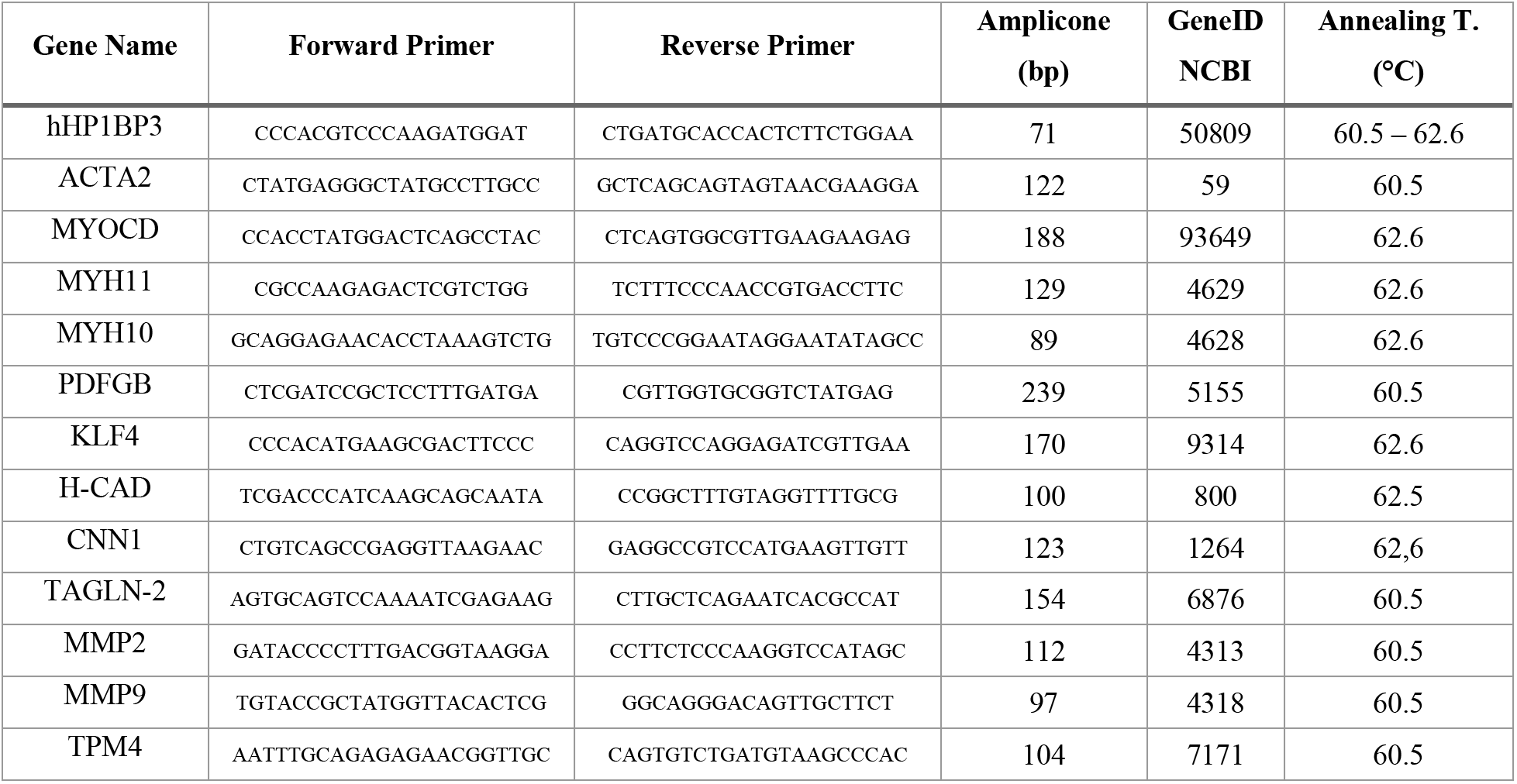
Primers used for qPCR.

**Table S5.** Antibodies used for flow cytometry.

| Primary Antibody | Fluor-chrome | Clone | D. F. | Cat. N. | Supplier |
| --- | --- | --- | --- | --- | --- |
| Zombie NIR | APC-eFluor780 | - | 1:200 | 423105 | Biolegend |
| CD14 | BV785 | HCD14 | 1:200 | 325628 | Biolegend |
| CD31 | BV605 | WM59 | 1:200 | 303121 | Biolegend |
| CD45 | PerCP/Cy5.5 | 2D1 | 1:200 | 368504 | Biolegend |
| Endoglin (CD105) | Pacific Blue - BV421 | SN6h | 1:200 | 800510 | Biolegend |
| VAP1 (AOC3) | APC -AF647 | 393106 | 1:200 | IC39571R | R&D System |
| CD144 | PE/Cy7 | BV9 | 1:200 | 348506 | Biolegend |
| ICAM-1 (CD54) | AF488 | HCD54 | 1:200 | 322713 | Biolegend |
| E-Selectin (CD62E) | PE/Cy7 | HAE-1f | 1:200 | 336015 | Biolegend |
| P-Selectin (CD62P) | BV785 | AK4 | 1:200 | 304941 | Biolegend |
| VCAM-1 (CD106) | BV421 | STA | 1:200 | 305815 | Biolegend |
| CD142 | PE | NY2 | 1:200 | 365203 | Biolegend |
| MCP-1 | APC | 5D3-F7 | 1:200 | 502611 | Biolegend |

## References

1. Alexander MR, Owens GK. Epigenetic control of smooth muscle cell differentiation and phenotypic switching in vascular development and disease. Annu Rev Physiol. 2012;74:13–40.

2. Martos-Rodríguez CJ, Albarrán-Juárez J, Morales-Cano D, Caballero A, MacGrogan D, de la Pompa JL, et al. Fibrous Caps in Atherosclerosis Form by Notch-Dependent Mechanisms Common to Arterial Media Development. Arterioscler Thromb Vasc Biol. 2021 Sep;41(9):e427–39.

3. Newman AAC, Serbulea V, Baylis RA, Shankman LS, Bradley X, Alencar GF, et al. Multiple cell types contribute to the atherosclerotic lesion fibrous cap by PDGFRβ and bioenergetic mechanisms. Nat Metab. 2021 Feb;3(2):166–81.

4. Hartman RJG, Owsiany K, Ma L, Koplev S, Hao K, Slenders L, et al. Sex-Stratified Gene Regulatory Networks Reveal Female Key Driver Genes of Atherosclerosis Involved in Smooth Muscle Cell Phenotype Switching. Circulation. 2021 Feb 16;143(7):713–26.

5. Wirka RC, Wagh D, Paik DT, Pjanic M, Nguyen T, Miller CL, et al. Atheroprotective roles of smooth muscle cell phenotypic modulation and the TCF21 disease gene as revealed by single-cell analysis. Nat Med. 2019 Aug;25(8):1280–9.

6. Owens GK, Kumar MS, Wamhoff BR. Molecular regulation of vascular smooth muscle cell differentiation in development and disease. Physiol Rev. 2004 Jul;84(3):767–801.

7. Shankman LS, Gomez D, Cherepanova OA, Salmon M, Alencar GF, Haskins RM, et al. KLF4 Dependent Phenotypic Modulation of SMCs Plays a Key Role in Atherosclerotic Plaque Pathogenesis. Nat Med. 2015 Jun;21(6):628–37.

8. Alencar GF, Owsiany KM, Karnewar S, Sukhavasi K, Mocci G, Nguyen AT, et al. Stem Cell Pluripotency Genes Klf4 and Oct4 Regulate Complex SMC Phenotypic Changes Critical in Late-Stage Atherosclerotic Lesion Pathogenesis. Circulation. 2020 Nov 24;142(21):2045–59.

9. Bennett MR, Evan GI, Schwartz SM. Apoptosis of human vascular smooth muscle cells derived from normal vessels and coronary atherosclerotic plaques. J Clin Invest. 1995 May;95(5):2266–74.

10. As D, Dk A. Inability of vascular smooth muscle cells to proceed beyond S phase of cell cycle, and increased apoptosis in symptomatic carotid artery disease. J Vasc Surg [Internet]. 2003 Jul [cited 2022 Jun 16];38(1). Available from: https://pubmed.ncbi.nlm.nih.gov/12844105/

11. Pankajakshan D, Jia G, Pipinos I, Tyndall SH, Agrawal DK. Neuropeptide Y receptors in carotid plaques of symptomatic and asymptomatic patients: effect of inflammatory cytokines. Exp Mol Pathol. 2011 Jun;90(3):280–6.

12. Koenen RR, Weber C. Chemokines: established and novel targets in atherosclerosis. EMBO Mol Med. 2011 Dec;3(12):713–25.

13. Bennett S, Breit SN. Variables in the isolation and culture of human monocytes that are of particular relevance to studies of HIV. J Leukoc Biol. 1994 Sep;56(3):236–40.

14. Novikova OA, Nazarkina ZK, Cherepanova AV, Laktionov PP, Chelobanov BP, Murashov IS, et al. Isolation, culturing and gene expression profiling of inner mass cells from stable and vulnerable carotid atherosclerotic plaques. PLoS ONE. 2019 Jun 26;14(6):e0218892.

15. Wang D, Wang Z, Zhang L, Wang Y. Roles of Cells from the Arterial Vessel Wall in Atherosclerosis. Mediators Inflamm. 2017 Jun 7;2017:e8135934.

16. Hu Z, Liu W, Hua X, Chen X, Chang Y, Hu Y, et al. Single-Cell Transcriptomic Atlas of Different Human Cardiac Arteries Identifies Cell Types Associated With Vascular Physiology. Arterioscler Thromb Vasc Biol. 2021 Apr;41(4):1408–27.

17. Visconti RP, Awgulewitsch A. Topographic patterns of vascular disease: HOX proteins as determining factors? World J Biol Chem. 2015 Aug 26;6(3):65–70.

18. Spanjersberg TCF, Oosterhoff LA, Kruitwagen HS, Dungen NAM van den, Harakalova M, Mokry M, et al. Locational memory of macrovessel vascular cells is transcriptionally imprinted [Internet]. bioRxiv; 2021 [cited 2022 Jul 7]. p. 2021.10.20.465092. Available from: https://www.biorxiv.org/content/10.1101/2021.10.20.465092v1

19. Depuydt MAC, Prange KHM, Slenders L, Örd T, Elbersen D, Boltjes A, et al. Microanatomy of the Human Atherosclerotic Plaque by Single-Cell Transcriptomics. Circ Res. 2020 Nov 6;127(11):1437–55.

20. Pan H, Xue C, Auerbach BJ, Fan J, Bashore AC, Cui J, et al. Single-Cell Genomics Reveals a Novel Cell State During Smooth Muscle Cell Phenotypic Switching and Potential Therapeutic Targets for Atherosclerosis in Mouse and Human. Circulation. 2020 Nov 24;142(21):2060–75.

21. Bennett MR, Sinha S, Owens GK. Vascular Smooth Muscle Cells in Atherosclerosis. Circ Res. 2016 Feb 19;118(4):692–702.

22. Gomez D, Shankman LS, Nguyen AT, Owens GK. Detection of histone modifications at specific gene loci in single cells in histological sections. Nat Methods. 2013 Feb;10(2):171–7.

23. Gomez D, Swiatlowska P, Owens GK. Epigenetic Control of Smooth Muscle Cell Identity and Lineage Memory. Arterioscler Thromb Vasc Biol. 2015 Dec;35(12):2508–16.

24. Pirillo A, Norata GD, Catapano AL. LOX-1, OxLDL, and atherosclerosis. Mediators Inflamm. 2013;2013:152786.

25. Pustlauk W, Westhoff TH, Claeys L, Roch T, Geißler S, Babel N. Induced osteogenic differentiation of human smooth muscle cells as a model of vascular calcification. Sci Rep. 2020 Apr 6;10(1):5951.

26. van Gils MJ, Vukadinovic D, van Dijk AC, Dippel DWJ, Niessen WJ, van der Lugt A. Carotid atherosclerotic plaque progression and change in plaque composition over time: a 5-year follow-up study using serial CT angiography. AJNR Am J Neuroradiol. 2012 Aug;33(7):1267–73.

27. Rohwedder I, Montanez E, Beckmann K, Bengtsson E, Dunér P, Nilsson J, et al. Plasma fibronectin deficiency impedes atherosclerosis progression and fibrous cap formation. EMBO Mol Med. 2012 Jul;4(7):564–76.

28. Moore KJ, Fisher EA. The double-edged sword of fibronectin in atherosclerosis. EMBO Mol Med. 2012 Jul;4(7):561–3.

29. Xie C, Ritchie RP, Huang H, Zhang J, Chen YE. Smooth muscle cell differentiation in vitro: models and underlying molecular mechanisms. Arterioscler Thromb Vasc Biol. 2011 Jul;31(7):1485–94.

30. Tabas I, García-Cardeña G, Owens GK. Recent insights into the cellular biology of atherosclerosis. J Cell Biol. 2015 Apr 13;209(1):13–22.

31. Feil S, Fehrenbacher B, Lukowski R, Essmann F, Schulze-Osthoff K, Schaller M, et al. Transdifferentiation of vascular smooth muscle cells to macrophage-like cells during atherogenesis. Circ Res. 2014 Sep 12;115(7):662–7.

32. Qiao JH, Mishra V, Fishbein MC, Sinha SK, Rajavashisth TB. Multinucleated giant cells in atherosclerotic plaques of human carotid arteries: Identification of osteoclast-like cells and their specific proteins in artery wall. Exp Mol Pathol. 2015 Dec 1;99(3):654–62.

33. Alexopoulos N, Raggi P. Calcification in atherosclerosis. Nat Rev Cardiol. 2009 Nov;6(11):681–8.

34. de Bakker M, Timmerman N, van Koeverden ID, de Kleijn DPV, de Borst GJ, Pasterkamp G, et al. The age-and sex-specific composition of atherosclerotic plaques in vascular surgery patients. Atherosclerosis. 2020 Oct;310:1–10.

35. Hashimshony T, Wagner F, Sher N, Yanai I. CEL-Seq: single-cell RNA-Seq by multiplexed linear amplification. Cell Rep. 2012 Sep 27;2(3):666–73.

36. Hashimshony T, Senderovich N, Avital G, Klochendler A, de Leeuw Y, Anavy L, et al. CEL-Seq2: sensitive highly-multiplexed single-cell RNA-Seq. Genome Biol. 2016 Apr 28;17(1):77.

37. Rosenberg AB, Roco CM, Muscat RA, Kuchina A, Sample P, Yao Z, et al. Single-cell profiling of the developing mouse brain and spinal cord with split-pool barcoding. Science. 2018 Apr 13;360(6385):176–82.

